# Reconciling Ecogeographical Rules: Rainfall and Temperature Predict Global Colour Variation in the Largest Bird Radiation

**DOI:** 10.1101/496463

**Authors:** Kaspar Delhey, James Dale, Mihai Valcu, Bart Kempenaers

## Abstract

Ecogeographical rules that associate climate with organismal form and function can reveal patterns of climatic adaptation. Two rules link animal coloration with climate: Gloger’s rule (darker coloration where wet and warm), and Bogert’s rule (darker coloration where cold). Whereas Gloger’s rule was proposed for endotherms, and Bogert’s rule for ectotherms, both rules may apply more broadly, despite their seemingly opposing effects. Here we test this contradiction on a global scale across passerine birds. Consistent with Gloger’s rule, birds were darker in wetter areas and, following Bogert’s rule, lighter where warm, although birds became lighter again at very low temperatures. Rainfall and temperature had antagonistic or additive effects depending on their pattern of covariation, and this predicted whether birds followed the rules. We integrate both rules into a general framework to explain heterogeneity in climatic effects on coloration, which has implications to understand patterns of diversification, climatic adaptation and climate change impacts.

## INTRODUCTION

Certain patterns of global spatial variation in organismal form and function are strong and general enough to have been formalised as ecogeographical rules (Gaston *et al*. 2008). However, the generality of these rules has been debated (Zink & Remsen 1986; Lomolino *et al*. 2006; Hagen 2017). Currently, even some of the best-studied ecogeographical rules, such as Bergmann’s rule (which links body size with temperature variation), are sources of controversy (Olson *et al*. 2009; Salewski & Watt 2017; Riemer *et al*. 2018). These debates are relevant today, because adaptive patterns of phenotypic variation linked to climate may help us predict potential responses to human-induced climate change (Millien *et al*. 2006; Roulin 2014; Teplitsky & Millien 2014). Hence, identifying the reasons why ecogeographical rules apply in some cases but not in others, is essential to establish their general validity and utility. However, our current knowledge is mainly restricted to body size variation, and this limits our understanding of the broad palette of adaptations to climate variation which include, among many others, appendage size, physiology, insulation, behaviour and coloration.

One of the oldest ecogeographical rules that links variation in animal coloration with climatic variation is Gloger’s rule (Delhey 2017). This rule was inspired by the work of CWL Gloger who recognised in 1833 that birds and mammals tend to be more intensively pigmented in tropical regions (Gloger 1833). In its simple modern interpretation, Gloger’s rule predicts that animals should be darker in warm and humid areas, such as tropical rainforests (Rensch 1929; Delhey 2017). Evidence supporting Gloger’s rule has been found in different groups of animals (Miskimen 1972; Zink & Remsen 1986; Lai *et al*. 2008, 2016; Roulin *et al*. 2009; Pannkuk *et al*. 2010; Friedman & Remeš 2017; Delhey 2018). However, because the tropics are associated with a variety of climatic factors, different patterns of variation have been considered consistent with Gloger’s rule, including correlations with rainfall, temperature, UV radiation, latitude, evapotranspiration or combinations thereof (Gipson *et al*. 2002; Caro 2005; Kamilar & Bradley 2011; Singaravelan *et al*. 2013; Bastide *et al*. 2014; Koski & Ashman 2015).

Indeed, recent comparative analyses on birds have revealed patterns that seemingly oppose some aspects of Gloger’s rule, namely that species with darker plumage are often found in colder rather than warmer regions (Friedman & Remeš 2017; Delhey 2018; Galván *et al*. 2018; Medina *et al*. 2018). These conflicting results have also been found by some studies of intraspecific variation (Aldrich & James 1991; Morales *et al*. 2017; Fargallo *et al*. 2018). However, darker animals occurring in colder regions is consistent with another ecogeographical rule: Bogert’s rule (Bogert 1949; Clusella Trullas *et al*. 2007; Gaston *et al*. 2009). Also called the thermal melanism hypothesis, this rule was originally proposed for ectothermic animals and argues that darker animals occur in colder regions for thermoregulatory reasons because dark coloration absorbs more solar radiation (Clusella Trullas *et al*. 2007).

Discrepancies between studies suggests that the correlation between climatic variables and animal light-to-dark variation differs across the globe. For example, the relationship between lightness and temperature could be positive in places such as Australia (Friedman & Remeš 2017; Delhey 2018) or Spain (Galván *et al*. 2018), where the need for thermoregulation to avoid overheating may play an important role, and negative at higher latitudes, driven by the need for camouflage against snowy surfaces. Thus, heterogeneous patterns of correlations in different biogeographic regions are not unexpected. An effective way to disentangle these possibilities is to assess the links between specific climatic variables and animal coloration on a global scale.

Resolving the nature and generality of correlations between climate and coloration is a first necessary step to identify the key drivers of ecogeographic rules. Positive correlations between animal lightness and temperature as suggested by Bogert’s rule are usually interpreted in terms of thermoregulation (Clusella Trullas *et al*. 2007), while the mechanisms behind Gloger’s rule are more elusive. Camouflage is widely considered a valid explanation for the occurrence of darker animals in more humid and vegetated environments, and of lighter animals in polar regions or deserts (Zink & Remsen 1986; Caro 2005), but other explanations exist (Burtt & Ichida 2004; Ducrest *et al*. 2008; Delhey 2017). If adaptive mechanisms underlie the patterns of geographic variation in coloration, then climate change and human-driven habitat transformation may disrupt these with potential negative consequences (Delhey & Peters 2017). For example, consistent with Bogert’s rule, insect assemblages have become lighter in regions experiencing increases in temperature, caused by shifts in distribution and local extinctions (Zeuss *et al*. 2014; Bishop *et al*. 2016). Historical changes in coloration across different species of birds and mammals have also been linked to climatic or environmental change (Galeotti *et al*. 2009; Maloney *et al*. 2009; Sandoval-Castellanos *et al*. 2017) and failure to adapt may result in local extinctions (Mills *et al*. 2013). Knowing the generality of climatic effects on colour variability is thus essential to help us understand and predict the effects of future changes.

Here we present a global test of both Gloger’s and Bogert’s rules by assessing the relationship between climatic (temperature and rainfall) and environmental variables (tree cover), and plumage lightness across nearly all species of passerine birds (Order Passeriformes).

## MATERIAL AND METHODS

### Plumage lightness

We quantified plumage lightness using Red-Green-Blue (RGB) values of plumage colours obtained from the plates of the Handbook of the Birds of the World (del Hoyo, J., Elliott, A. & Christie n.d.) as described in Dale et al. (Dale *et al*. 2015) for all species of the order Passeriformes (n = 5831). Briefly, we obtained RGB values from scanned plates for nine different plumage patches (nape, crown, forehead, throat, upper breast, lower breast, belly, vent and back). Each separate channel (R, G, and B) can vary between 0 and 255, and we compute plumage lightness as (R+G+B)/3, which therefore also varies between 0 (pure black) and 255 (pure white, see Supplementary Information and Figs. S1-S2). For each species, we averaged these values across all patches, but separately for each sex, to obtain sex-specific estimates of plumage lightness. Male and female lightness scores were positively correlated (r = 0.8, 95%CI = 0.79 – 0.81, t = 102.3, df = 5807, p < 0.001). As shown in other studies using reflectance spectrometry (Delhey 2018) or book plates (Negro *et al*. 2018), males are usually darker than females (mean differences within species [female lightness – male lightness] = 8.3, 95%CI = 7.7 – 8.8, paired t=28.5, df=5808, p < 0.001).

The lightness estimates used in this study did not cover the entire body, and were derived from book plates rather than from plumage reflectance spectra. Hence, we assessed how well they correlated with spectrometric measurements of plumage lightness derived from the entire plumage of museum specimens (Delhey 2018) for the subsample of species for which we had both types of measurements (n = 309). For both sexes, RGB values correlated with measurements of real plumage lightness (males: r = 0.72, females: r = 0.70, both p < 0.001). To further determine whether differences between methods could affect our conclusions, we tested whether climatic effects were consistent between both methods (for the subsample of 309 species). This was the case; both conclusions and effects are similar (Fig. S3). Hence, we consider the RGB values used here to constitute an appropriate estimate of overall plumage lightness.

### Climatic variables and vegetation

Based on the breeding range of each species from (Dale *et al*. 2015) we obtained species-specific average estimates of mean annual temperature (bio1, in °C) and annual average precipitation (bio12, in mm, square-root transformed) from the database in (Karger *et al*. 2017) and percent forest cover from (DeFries *et al*. 2000), sampled on a grid of 0.05° cell size using the package ‘RangeMapper’ (Valcu *et al*. 2012). Similarly, for assemblage level analysis we obtained average estimates for annual mean temperature, annual rainfall (square-root transformed) and forest cover within each 1° cell (∼100 x 100 km).

### Statistical Analyses

#### Analysis at species level

We used the function phylolm from the R package ‘phylolm’ (Tung Ho & Ané 2014) with plumage lightness as the dependent variable and temperature, rainfall and percent tree cover as independent variables. We also included the quadratic effects for precipitation, temperature and tree cover to assess for possible non-linear effects. All variables were centred and scaled prior to analyses to facilitate the interpretation of linear and quadratic effects (Schielzeth 2010). We controlled for phylogenetic relatedness using Pagel’s lambda model. A sample of 1000 phylogenetic trees for the species included was obtained from birdtree.org (Jetz *et al*. 2012). Models were run on the 1000 phylogenies and we obtained average effect sizes and their SEs using model averaging procedures (Symonds & Moussalli 2011) across all models. In addition, we also computed median, minimum, and maximum p-values, as well as the proportion of phylogenies yielding a p-value < 0.05 for each parameter. Analyses were carried out separately for male and female lightness. Complete environmental and phylogenetic data were available for 5809 out of a total of 5831 species for which we had colour data (99.6%). Since the function phylolm does not include R^2^-values in its output, we assessed fit by correlating predicted and observed lightness values. To determine the extent of taxonomic heterogeneity in the relationships between plumage lightness and rainfall, temperature and tree cover, we ran the comparative analyses separately for each family comprising 10 or more species. However, in this case, we used only a sample of 100 phylogenies. Inspection of model residuals did not reveal major departures from normality or heterogeneity of variance.

#### Analysis at assemblage level

For each 1° cell we computed a mean plumage lightness value for all species in the assemblage, separately for males and females. Only cells with more than 5 species were included, yielding a total of 9715 cells. Climatic variables and tree cover were also summarised in the same manner. Mean assemblage lightness was the dependent variable, while precipitation, temperature and percent tree cover were included as explanatory variables (as linear and quadratic effects). All variables were centred and scaled prior to analyses (Schielzeth 2010). We also included zoogeographical realm (11 levels, (Holt *et al*. 2013)) as a random intercept to account for differences in lightness across the globe. These zoogeographic realms define terrestrial areas that harbour subsets of species that are closely related, and which are often part of clades that have radiated in, and are restricted to, specific regions of the globe. Terrestrial realms therefore constitute units representing largely independent macro-ecological and -evolutionary patterns. We further included random-slope effects for temperature, precipitation and percent tree cover (linear and quadratic where relevant) in each realm to allow the possibility of different patterns in each realm. We accounted for spatial autocorrelation of residuals based on the approach outlined by Crase et al. (Crase *et al*. 2012) which involves fitting a residual autocorrelation covariate derived from a 5×5 cell focal region around each cell. This procedure reduced spatial autocorrelation as revealed by visual inspection of variograms. Model residuals did not show major departures from normality or heterogeneity of variance. We computed marginal (fixed effects) and conditional (mixed+fixed effects) R^2^-values following (Nakagawa & Schielzeth 2013) using the package MUMin (Barton 2017). Because the spatial autocorrelation covariate accounts for much of the unexplained variation (and thus increases R^2^-values beyond the contribution of the other fixed effects), R^2^-values were computed on the model without this variable. Assemblage-level analyses were carried out with the R package lme4 (Bates *et al*. 2015).

#### Simulations

Assemblage level analyses that seek to link climatic variables with trait data and use averages for each assemblage, suffer from a high probability of type I errors, even after accounting for spatial autocorrelation (Friedman & Remeš 2017; Delhey 2018). To assess whether this affected our conclusions we simulated the evolution of plumage lightness values for each species across the phylogeny assuming Brownian motion for each of the 1000 phylogenies used in the species-level analyses. Simulations were carried out using the function ‘fastBM’ from the package ‘phytools’ (0.6-44) (Revell 2012), separately for female and male lightness. For each phylogeny we first quantified the parameters λ and σ^2^ based on the real lightness values, and we then used these parameters to simulate lightness values. Given that these simulated values of species lightness should only be influenced by phylogenetic relatedness, we would not expect any correlation with climatic variables beyond those that are inherent when related species share climatic niches. Based on the simulated data we re-ran the final best model for the assemblage-level analyses 1000 times to obtain a distribution of simulated effects. We then compared the observed effects from the real data with the simulated data, and computed the proportion of simulated effects that were larger than the observed effect.

#### Meta-regressions

We used a meta-regression approach to determine whether variability in the patterns of precipitation-temperature covariance could explain variability in climatic or tree cover effects across bird families or zoogeographic realms. A meta-regression is a meta-analysis that includes the effect of a continuous covariate (also called moderator (Nakagawa & Santos 2012)). Meta-analyses combine statistical effects from different analyses, while accounting for their measurement error, to obtain an average effect across analyses, and test for variables that explain differences between analyses. We ran two groups of meta-regressions, one using each family-specific effect of precipitation, temperature or tree cover on plumage lightness, as derived from linear models accounting for phylogenetic relatedness (see species-level analyses above). We used only families with at least 10 species but results are comparable if we only used families with at least 15 species (see Results). For the second group of meta-regressions we used each zoogeographic realm specific effect, while controlling for spatial effects (see assemblage-level analyses above). In both cases we used the function ‘rma’ (method=’REML’) in the package metafor (Viechtbauer 2010) to run the meta-regressions, which weigh each effect by its variance.

## RESULTS

Birds living closer to the equator have darker plumage (Figs. 1-2). This pattern was consistent at both the species level (latitude effect: estimate females = 0.173, SE = 0.015, median p < 0.001; estimate males = 0.128, SE = 0.015, median p < 0.001), and the assemblage level (latitude effect: female estimate = 0.398, SE = 0.003, p < 0.001; male estimate = 0.573, SE = 0.003, p < 0.001). At the assemblage level, plumage became darker at higher latitudes (Fig. 1, quadratic latitude effect: female estimate = −0.414, SE = 0.003, p < 0.001, male estimate = −0.361, SE = 0.003, p < 0.001), but this was not as evident at the species level (Fig. 1, quadratic latitude effect: female estimate = - 0.009, SE = 0.000, median p = 0.316, male estimate = −0.011, SE = 0.009, median p = 0.215).

**Figure 1.**
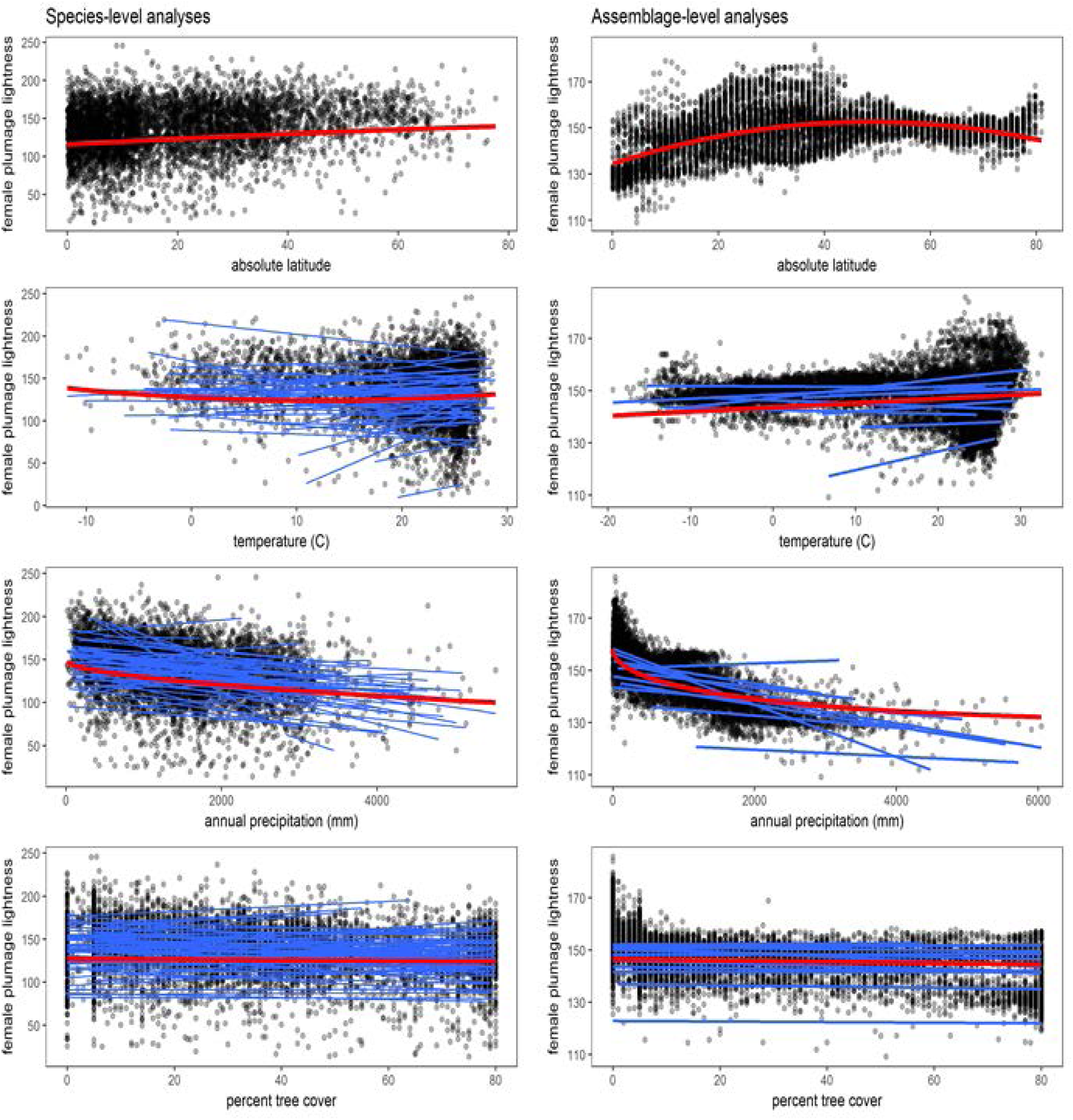
Correlates of plumage lightness in female passerines. Scatterplots depict female plumage lightness for all species of passerine birds in relation to different geographic (latitude), climatic (annual mean temperature, annual precipitation) and environmental (percent forest cover) predictors at the species level (A) – D), n = 5809) and assemblage level (E) – H), n = 9715). Red lines depict the global line of best fit derived from the best models (except for latitude, which was fitted only to show broad patterns of variation); blue lines depict lines of best fit (linear effects only) derived separately for each family (species level analyses, n = 65 families with 10 or more species each) or zoogeographic realms (assemblage level analyses, 11 realms). Note that while scatterplots of temperature effects suggest heterogeneity of variance, this was not evident in the model residuals.

**Figure 2.**
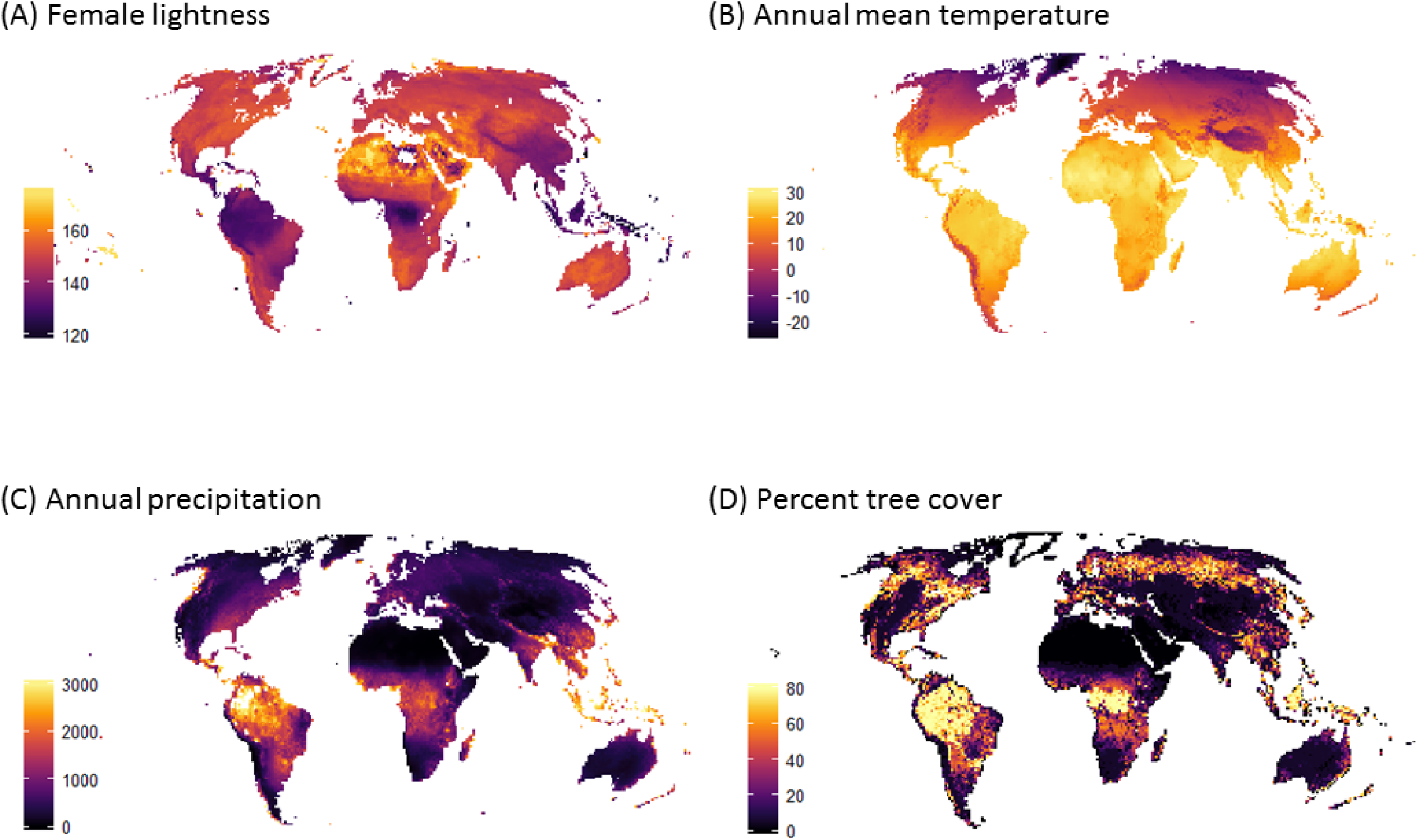
Maps illustrating the geographic distribution of (A) mean female plumage lightness, (B) annual mean temperature, (C) annual precipitation and (D) percent tree cover.

At the species level, the main correlate of plumage lightness was annual precipitation (Figs. 1, 3). Species living in areas with high levels of precipitation and tree cover were darker (Figs. 1, 3). The quadratic precipitation and tree cover effects were never significant and excluded from the models (Fig. 3). The quadratic temperature effect was positive and statistically significant indicating that lightness was lowest at intermediate temperatures (∼10 °C annual mean temperature, Fig. 1, 3). The overall linear effect of temperature was positive (Fig. 3). In general, effects were stronger for female plumage (model R^2^ = 0.11) than for male plumage (model R^2^ = 0.06), but patterns were similar (Fig. 3, for full statistical results see Table S1). Separate analyses for each bird family showed that the effects of precipitation, tree cover and temperature can vary substantially among families (Fig. 1).

**Fig. 3.**
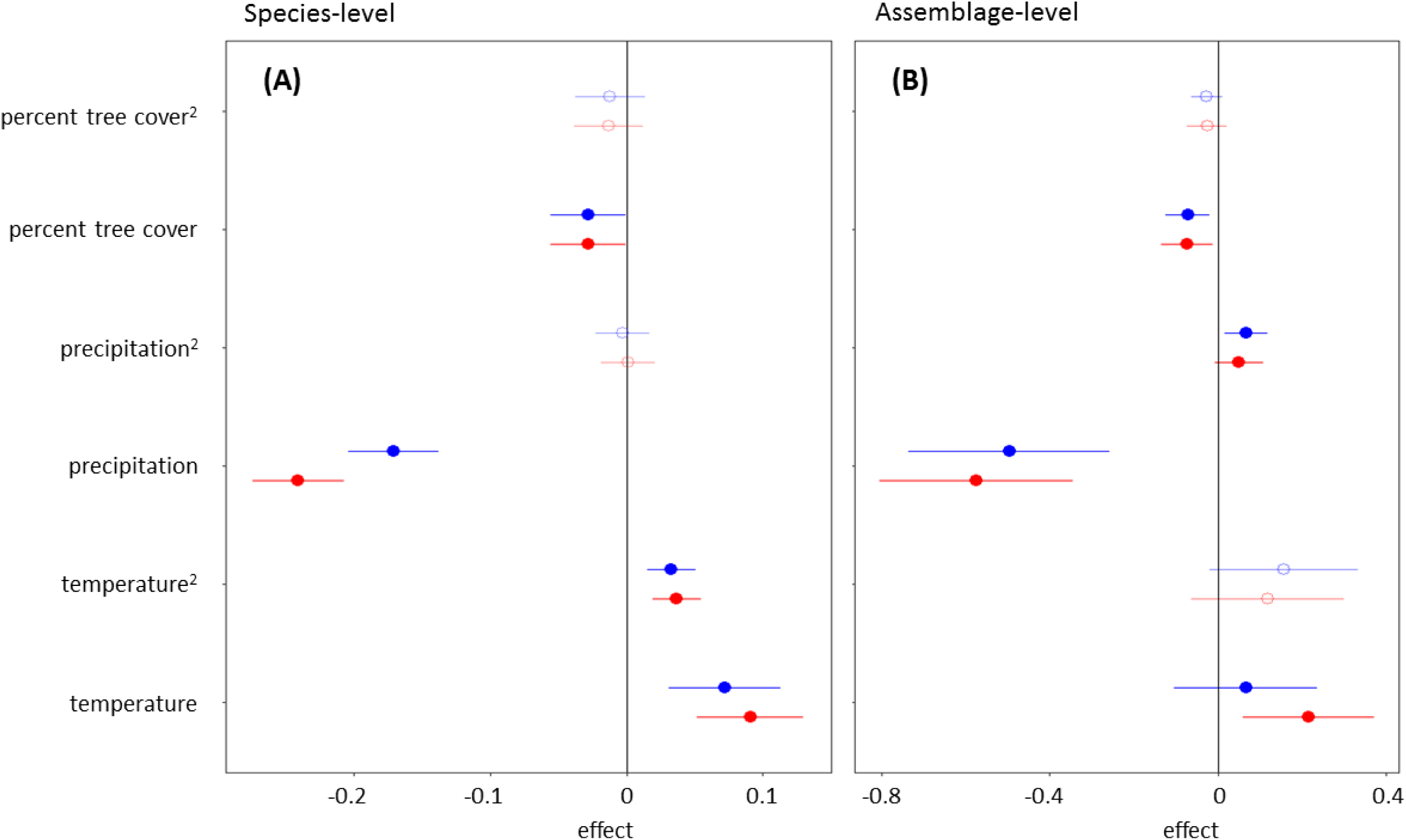
Predictors of plumage lightness in passerine birds. Forest plots depict effects (±95% CI) of each predictor on plumage lightness of females (red symbols) and males (blue symbols) for analyses at the species (A) and assemblage (B) levels. Non-significant quadratic effects excluded from the final model are depicted by open symbols, with the exception of the quadratic precipitation effect at the assemblage level which was not significant in females but significant in males. Positive effects represent increases in plumage lightness with increases of the explanatory variable. All variables are scaled, hence each effect represents the change in the response variable (in standard deviations) associated with a change in one standard deviation of the explanatory variable. For full statistical results see Tables S1-S2.

At the assemblage level, bird communities in areas with higher levels of precipitation and tree cover contained, on average, species with darker plumage (Figs. 1-3). While the quadratic tree cover effect was not significant (and therefore excluded), the model revealed a significant quadratic precipitation effect in males and the same trend in females (Fig. 1), indicating that the precipitation effect weakens at very high levels of precipitation (Fig. 1). Linear temperature effects were largely positive (but not statistically significant in males), with lighter assemblages occurring in warmer areas (Figs. 1-2); no significant quadratic temperature effects were detected (Fig. 3). Fixed effects alone explained a substantial amount of variation in assemblage lightness (marginal R^2^_females_ = 0.32; marginal R^2^_males_ = 0.25); when random effects were taken into account the amount of variation explained was even higher (conditional R^2^_females_ = 0.78, conditional R^2^_males_ = 0.75). For full statistical results see Table S2.

Since assemblage-level analyses can be prone to inflated type I error rates (Friedman & Remeš 2017; Hawkins *et al*. 2017; Delhey 2018), we compared the observed results with those obtained from a sample of 1000 phylogenetic simulations (see methods). This approach indicated that our conclusions are robust: observed effects in the final assemblage model (Fig. 3) were always larger than the most extreme simulated effects.

Across zoogeographic realms (Holt *et al*. 2013), there was substantial variation in average plumage lightness and the magnitude and direction of the temperature effect (Fig. 1, and as indicated by random intercept and slope variances, Table S2). Separate plots for each realm confirmed this, showing in particular negative effects of temperature for the Neotropical and Sino-Japanese realms (Fig. S4). Further exploratory plotting (Fig. S5) revealed that these realms showed some of the strongest positive correlations between annual precipitation and average temperature.

To evaluate this further, we tested whether variation in the correlation between temperature and precipitation could explain heterogeneity in climatic effects detected at the family and realm levels. If both climatic variables correlate in a strong positive manner, temperature and rainfall effects may oppose each other, weakening their correlation with plumage lightness. We used a meta-regression approach to test whether the effects of temperature, precipitation and tree cover – estimated separately for each family or realm and weighted by their uncertainty – co-varied with the strength of the temperature-rainfall correlation. For these analyses we only considered models with linear effects, rather than also including the quadratic effects. The main aim of these tests was to determine whether the overall positive or negative effects of temperature, precipitation and tree cover can be affected by antagonistic or additive effects of temperature and precipitation. Including quadratic effects makes interpreting these linear effects more difficult. Moreover, quadratic effects were statistically significant in only 5 families of birds.

At the species level, across passerine families, the positive effect of temperature on plumage lightness became significantly weaker as the correlations between precipitation and temperature became more positive (Table S3, Fig. 4a). Similarly, the negative effect of tree cover on plumage lightness became less marked with stronger positive temperature-precipitation correlation values (Table S3, Fig. 4c). The precipitation effect was not as affected by this correlation (Table S3, Fig. 4b). Results for males followed similar patterns, but effects were weaker (Table S3). It could be argued that the reason why effects become weaker when precipitation and temperature are highly correlated is simply due to the statistical effect of collinearity (F. Dormann *et al*. 2007). However, the fact that patterns are similar and in some cases became stronger if we tested for effects of each variable separately (thereby avoiding collinearity issues) (Table S3) argues against this.

**Figure 4.**
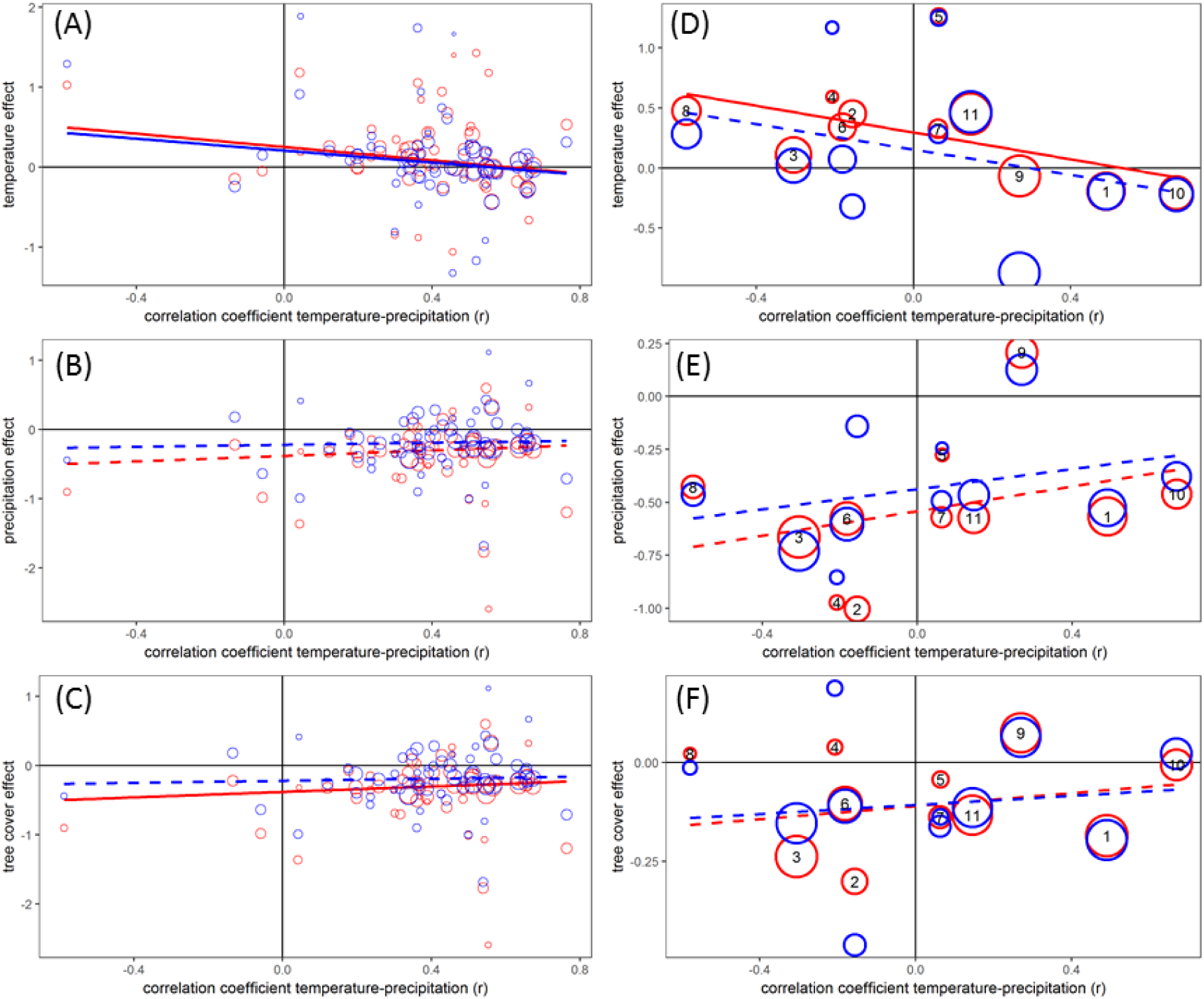
The association between the strength of the climatic (temperature [A, D], annual precipitation [B, E]) and environmental (tree cover [C, F]) effects on female (red) and male (blue) plumage lightness and the correlation between temperature and rainfall for different bird families (A) - C), 65 families with 10 or more species, but results are similar using only families with 15 or more species, Table S3) or zoogeographic realm (D – F), 11 realms). Each dot depicts a family- or realm-specific effect. Dot size is proportional to 1/SE of the effect (but on a different scale in A-C versus D-F). The lines show the best fit derived from meta-regression analyses weighing each effect by its uncertainty, continuous lines represent statistically significant effects (p < 0.05, for full results see Tables S3-S4). Numbers in D-F correspond to zoogeographic realms: 1: Neotropical, 2: Australian, 3: Aftrotropical, 4: Madagascan, 5: Oceania, 6: Oriental, 7: Panamanian, 8: Saharo-Arabian, 9: Nearctic, 10: Sino-Japanese, 11: Palearctic.

At the assemblage level, the effect of temperature on plumage lightness decreased with strong and positive temperature-precipitation correlations across the 11 zoogeographic realms (Table S4, Fig. 4d), while the effect of precipitation and tree cover became less marked (Table S4, Fig. 4ef). As above, results are qualitatively similar but weaker for males (Fig. 4d-f, Table S4), but again become more marked when we run models for each predictor separately (Table S4).

## DISCUSSION

Our global analysis is broadly supportive of the observations made by Gloger nearly 200 years ago: birds tend to be darker towards the equator (Gloger 1833). Assessing the effect of three possible environmental drivers behind this correlation reveals that annual precipitation is the most consistent correlate of plumage lightness: both at the species and assemblage levels, passerine birds are darker in areas with higher rainfall. Tree cover had a similar but weaker effect with darker birds inhabiting areas with higher tree cover. The effect of temperature was less strong, but in general birds tended to be lighter, not darker in warmer areas. Note, however, that at the species (but not assemblage) level, lightness increased again at very low temperatures. In sum, precipitation effects followed predictions by Gloger’s rule while temperature effects largely contradicted it, and were more consistent with Bogert’s rule.

These global patterns are in general agreement with recent work on Australian birds both at the species and assemblage levels, which showed similar negative precipitation effects and positive temperature effects (Friedman & Remeš 2017; Dalrymple *et al*. 2018; Delhey 2018). However, some of these effects vary in strength depending on the specific region (Dalrymple *et al*. 2018; Delhey 2018) or taxonomic group assessed (Friedman & Remeš 2017). Similarly, work on Spanish birds also reported a positive correlation between plumage lightness and temperature, but not with rainfall (Galván *et al*. 2018). This variability in climatic effects is reflected by our own analysis because the strength and direction of climatic effects varied across bird families and zoogeographic realms (Figs. 1, 4).

Our observations suggest that spatial and taxonomic heterogeneity of climatic effects can be partly explained by patterns of covariation between rainfall and temperature. When the correlation is strong and positive, the effects of rainfall and temperature on plumage lightness weaken (Fig. 4), because both gradients oppose each other (lightness correlates mostly positively with temperature and negatively with precipitation). This effect is more evident on temperature effects, which are generally much weaker than precipitation effects. Antagonistic climatic effects could provide a general explanation for a lack of agreement between studies. Trade-offs between different selective forces (which may or may not be summarised as ecogeographical rules) are not un-common, and have been used to explain, for example, departures from Bergmann’s rule (Medina *et al*. 2007; Gutiérrez-Pinto *et al*. 2014; De Keyser *et al*. 2015; Busso & Blanckenhorn 2018). However, previous studies have not explicitly identified the antagonistic or reinforcing effects of different climatic variables. We note that because we generated this explanation based on the data, proper confirmation will require further tests in different taxonomic groups and/or at a different level of variation (e.g. intraspecific variation).

The effects of temperature and precipitation are also constrained by the shape of the climatic space. This space, as defined by precipitation and temperature, is triangular in shape (Fig. 5), because very high levels of precipitation are always found in warm areas, while very cold regions always have low levels of precipitation. Very high levels of precipitation never coincide with very low temperatures. The consequence of this is that patterns of colour variation seem to follow the predictions of Gloger’s rule: birds are darker where it is warm and wet (and to some extent lighter where very cold). Yet, when assessing the underlying effects of the climatic variables, it becomes evident that this is only because at high temperatures the precipitation effects overwhelm temperature effects. Birds are dark despite high temperatures, not because of them. On the other hand, the lightening of plumage at cold temperatures (<10°C) does match the prediction of Gloger’s rule of lighter ‘polar coloration’ (Rensch 1929). We note, however, that this effect was only detected at the species-level analysis and that it applies to comparatively few species (∼10%); most passerines inhabit warmer areas, where temperature effects lead to lighter plumage (Fig. 5).

**Figure 5.**
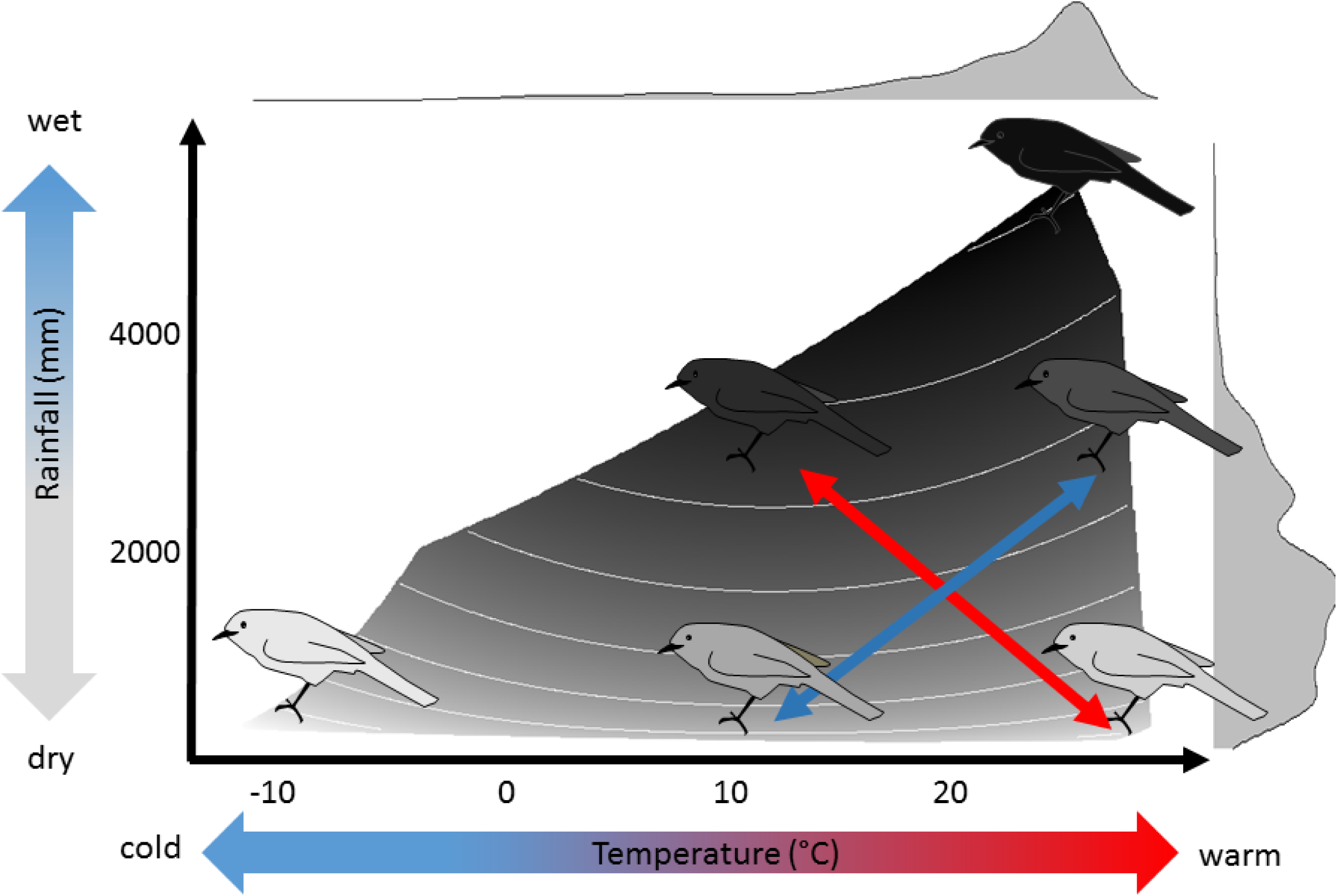
Proposed framework integrating the effects of precipitation (Gloger’s rule) and temperature (Bogert’s rule) on animal light-to-dark variation. The shades of the background and bird icons are based on predictions from the species-level model (Fig. 3A), which are restricted to the range of temperatures and precipitations present in the sample. The temperature effect shows a quadratic relationship: at high and low temperatures, plumage is lighter, at intermediate levels plumage is darker. Marginal density plots depict where in climatic space most species are found. Note that the lightening of plumage at low temperatures applies to comparatively few species because most passerines are found at higher temperatures where lightness increases with temperature. The precipitation effect (darker plumage at higher precipitation) is much stronger than any temperature effects in birds and this is why in warm and humid places birds are expected to have darker plumage. The two arrows depict two hypothetical climatic gradients. The red one is a gradient that should lead to high levels of colour divergence because temperature and precipitation effects work together (note how the arrow crosses 5 contour lines). The blue gradient leads to low levels of colour divergence because both gradients are working against each other (the arrow crosses only 3 contour lines).

Because Gloger’s rule generally predicts darker animals in warmer areas, and our temperature effects do not seem to follow this expectation in general, it is tempting to reformulate the rule in terms of precipitation alone rather than precipitation and temperature together. We therefore propose a general framework to explain climatic effects on plumage coloration that integrates and reconciles Gloger’s and Bogert’s rules, by taking into account their reinforcing or antagonistic effects (Fig. 5). Within this framework, for most species of passerines (and at the assemblage level) temperature effects on coloration seem to fall in the realm of Bogert’s rule. Gloger’s rule on the other hand, could be re-defined in terms of precipitation effects alone.

Traditionally, Bogert’s rule has been applied mainly to ectotherms (Clusella Trullas *et al*. 2007) and Gloger’s rule was formulated for endotherms, although the possibility that it also applies to ectotherms was left open (Rensch 1929). Past and recent work indicates that these rules could apply more broadly across animals (Amtmann 1965; Xing *et al*. 2016; Friedman & Remeš 2017; Cheng *et al*. 2018; Delhey 2018; Galván *et al*. 2018), fungi (Cordero *et al*. 2018) and perhaps even in plants (Watson & Hanham 1977; Burns 2015; Cuthill 2015). Whether patterns follow more strongly Gloger’s or Bogert’s rule could depend on the selective pressures behind the mechanisms in each rule. For example, for ectotherms thermoregulation may be more important such that correlations between lightness and temperature would be more prevalent. In contrast, for endotherms camouflage or parasite resistance (two popular adaptive explanations for Gloger’s rule (Delhey 2017)) could exert stronger selection, such that correlations with rainfall become more marked (as seems the case in birds).

This integrated framework has implications for further research. If climatic gradients where precipitation and temperature effects oppose each other lead to reduced colour divergence, pre-mating hybridization barriers may be reduced (Martin *et al*. 2010), and this could affect the maintenance of incipient species in cases of secondary sympatry. Hence, not only the magnitude (Jezkova & Wiens 2018) but also the nature of the climatic gradient may influence speciation in organisms where visual signals are important pre-mating barriers between species (e.g. birds). For example, together with song, differences in plumage coloration between subspecies of Song Sparrow (*Melospiza melodia*) across a rainfall gradient contribute to the prevention of hybrid mating (Patten *et al*. 2004). Hence, coloration could be considered as a “magic trait” (traits under divergent selection that also result in non-random mating (Servedio *et al*. 2011)), being affected by climatic conditions and playing a role in mate choice (Keller & Seehausen 2012). To our knowledge, no studies have linked lineage diversification patterns with the nature of climatic gradients. We predict that climatic gradients where rainfall and temperature effects reinforce each other (that is, negative correlation between rainfall and temperature) should be associated with higher rates of lineage diversification in animals that strongly rely on visual cues.

Furthermore, if patterns of covariation between climatic variables constrain the evolution of coloration that is optimally suited for the prevailing temperature or rainfall levels, alternative adaptations may evolve in compensation. These could include adaptations for thermoregulation such as insulation properties of fur or plumage (Briscoe *et al*. 2015), physiology (Tattersall *et al*. 2012), body and appendage size (Symonds & Tattersall 2010), or anti-predator behaviour to compensate for imperfect camouflage (Mcqueen *et al*. 2017). It could be argued that some of these compensatory responses should be evident in males (e.g. (Briscoe *et al*. 2015)), because male plumage lightness is less closely linked to climatic variation than female lightness. Males are under stronger sexual selection than females (Dale *et al*. 2015) and this often selects for conspicuous colours that function best when they stand out rather than blend in with the background. For example, plumage signals are selected to be lighter in environments with dense vegetation (Marchetti 1993), opposing the effects of Gloger’s rule described here. The fact that plumage coloration is a complex trait – the compromise of a variety of selection forces and evolutionary constraints – probably accounts for the generally modest explanatory power of climatic effects (Fig. 1).

In general, compensatory mechanisms that mediate evolutionary compromises may provide clues about possible alternative adaptive responses if future climate change disrupts current patterns of climatic covariation (Wu *et al*. 2011). More broadly, assessing the simultaneous effects of climate on a suite of adaptations can –not only help to establish why ecogeographical rules are not always followed– but also provide a more complete understanding on the diversity of phenotypic responses to climatic variation across taxa. To this end we need to expand the type of integrative framework used here to encompass multiple ecogeographic rules.

In conclusion, we propose a general framework to understand the effects of temperature and precipitation on the evolution of animal colours. Our data suggest that climatic effects on plumage coloration are affected by the type of climatic gradient, and that the strength and direction of climatic effects on plumage lightness can also vary across climatic space, while being constrained by the specific shape of the climatic space (Fig. 5). Given that the patterns uncovered at the species and assemblage levels should reflect selection at the individual level, studies on intraspecific variation in coloration are needed to put these predictions to test. Here we have deliberately focused on the two climatic variables that are most closely linked to the original ecogeographical rules. However, other climatic (and non-climatic) variables may be playing additional roles (Dalrymple *et al*. 2018) and correlations with temperature and precipitation may be driven by other causes. For example, it has been suggested that Gloger’s rule should be reformulated in terms of vegetation rather than precipitation because the former is presumably more closely linked to the putative mechanism (darker animals more camouflaged in darker environments)(Zink & Remsen 1986). This assumes that the mechanism behind Gloger’s rule is clear – an assumption that remains disputed (Burtt & Ichida 2004; Ducrest *et al*. 2008; Delhey 2017). More recently, research on the genetic basics melanin pigmentation –the main pigment involved in Gloger’s (Delhey 2017) and Bogert’s rules (Clusella Trullas *et al*. 2007), and the most common pigment in birds (Delhey 2015; Galván & Solano 2016)– has revealed pleiotropic links between genes that underlie pigmentation and multiple adaptive processes such as dispersal, aggressiveness, immune-responsiveness, stress-physiology, (Ducrest *et al*. 2008; San-jose & Roulin 2018) and even thermoregulation behaviours (Dreiss *et al*. 2016). Such links suggest that correlations between coloration and climate could be the consequence of selection that is not acting directly on colour per se. Hence, determining the mechanism(s) responsible for the correlations between climate and coloration may lead to further refinement and reformulation of the rule(s). In the meantime, our results provide an integrated framework to assess the generality of climatic effects on animal coloration, a necessary first step to determine their status as biological rules.

## Supporting information

## ACKNOWLEDGMENTS

We thank three anonymous reviewers for constructive criticism that helped improve the manuscript, in particular the suggestion to explore non-linear climatic effects. J.D. was supported by a Marsden Fund Grant (15-MAU-136) from the Royal Society of New Zealand.

