## Supplementary material for "Reconciling Ecogeographical Rules: Rainfall and Temperature Predict Global Colour Variation in the Largest Bird Radiation"

**Computing lightness estimates using RGB values**

In the RGB colour space, each channel (R, G or B) varies between 0 and 255. When all three channels have high values (i.e. close to 255) we perceive these colours as white, and when all three are low and close to 0 we experience these colours as black. In the figure below (Fig. S1) the RGB square, where each data point corresponds to an RGB value for the plumage patches measured in this study) high values for all RGB occur at the top right corner and low values for all RGB at the bottom left. Each symbol is coloured to resemble the original measured colours.

**Fig. S1.** Plumage colours of passerine birds plotted in RGB space.

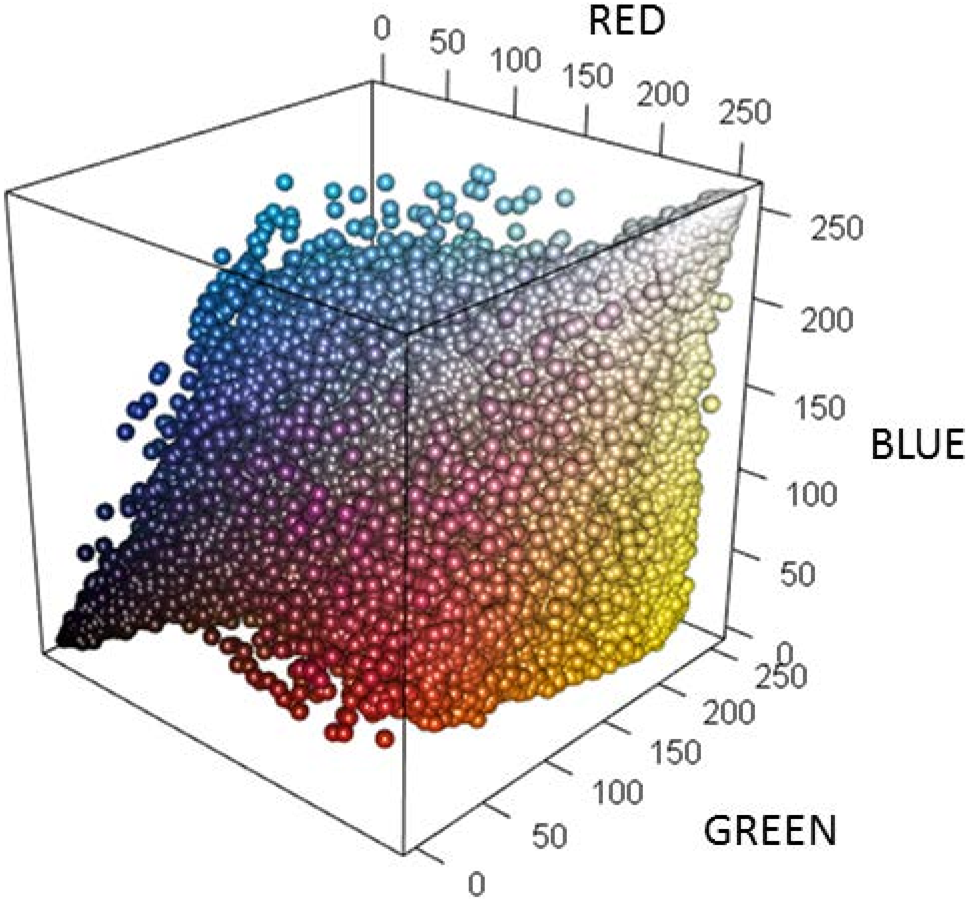


There are other colours spaces, which define lightness in slightly different ways. Converting RGB values into the HSL space (using function rgb2hsl from the package ‘plotwidgets’, (Weiner 2016)) and comparing the RGB lightness estimate used here with the HSL lightness estimate (‘L’) indicates that they are highly correlated (r = 0.99). Alternatively, we can assess whether there is a major axis of variation with similar positive loadings by R, G and B that reveals a common lightness dimension. If we run a principal component analysis, the first principal component accounts for 83% of the variation in RGB values and the RGB loadings are similar for all three channels (R = 0.58, G = 0.59, B = 0.55). More importantly, if we correlate this first principal component with our RGB lightness estimate, the correlation is nearly perfect (r = 0.999).

Finally, we can also compare well-known species at both ends of the spectrum, for example, a dark plumaged crow (*Corvus corone*) has an overall lightness estimate of 68, while the Bali Myna (*Leucopsar rothschildi*) - with its almost entirely white body plumage - has a value of 245. We can also plot the colours of each measured plumage patch against lightness, and again whites and light greys have high values of lightness, while blacks and dark greys have low values and the other colours fall in-between (Fig. S2). These tests and observations indicate that our lightness estimate should be well-suited at capturing the dark dark-to to-light variation that we are interested in.

**Fig. S2.** Plumage colours of passerine birds plotted against their RGB lightness estimates separately for females and males. Each point is coloured to resemble measured colours from the plates of the Handbook of the Birds of the World.

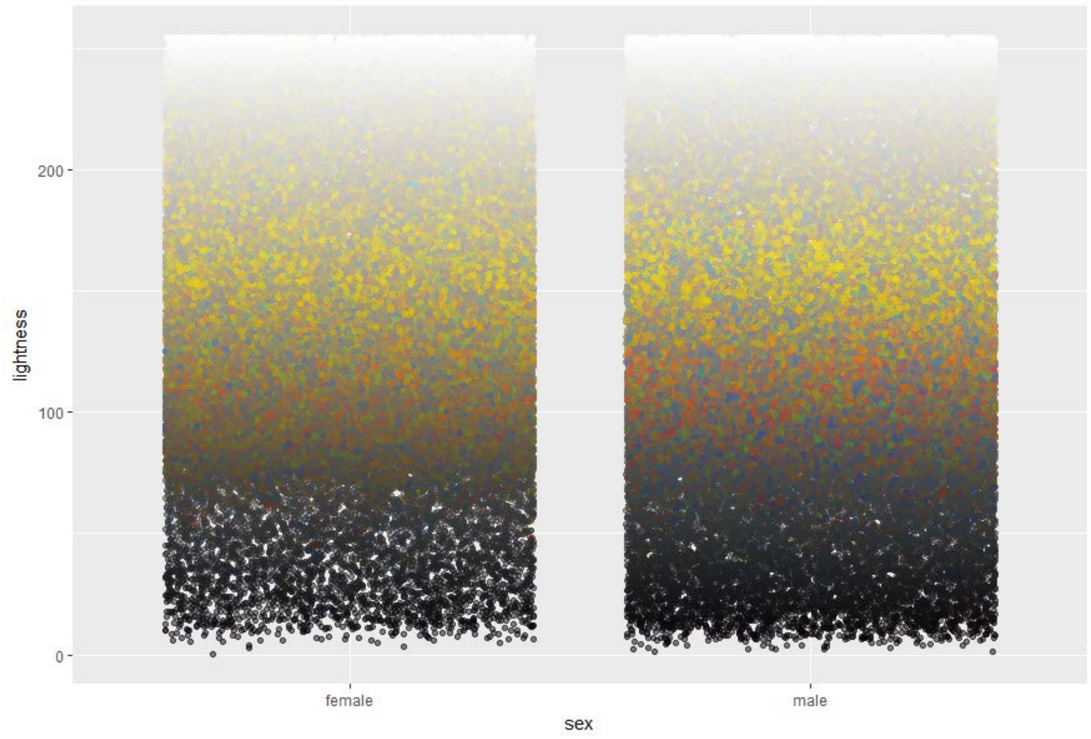

**Fig. S3.** Forest plot comparing temperature, precipitation and percent tree cover effect on plumage lightness using RGB lightness values or reflectance measurements (from (Delhey 2018)) as response variables. Forest plots depict effects (±95% CI) of each predictor on plumage lightness of females (red symbols) and males (blue symbols) for analyses at the species level (N = 309 species of Australian passerines). Note that in this case the effects of tree cover where assessed in a separate model since tree cover correlates strongly with precipitation (r = 0.8) in Australia which causes problems of collinearity, and that precipitation has been log10-transformed due to the extreme skew of this variable in Australia, see (Delhey 2018). Note as well that quadratic effects were excluded from this model as they were not significant (again similar to (Delhey 2018)).

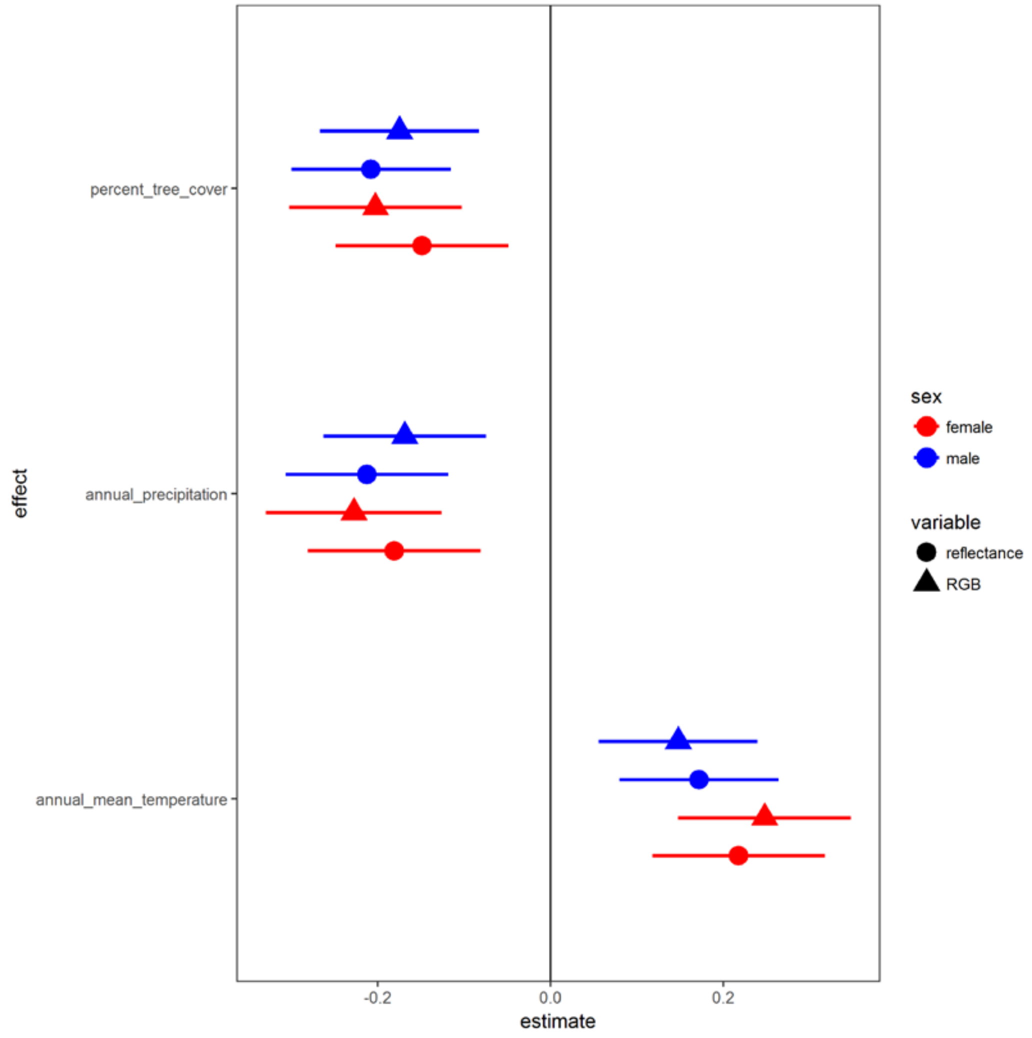

**Fig. S4.** Scatterplots depicting the correlation between annual mean temperature and average female lightness at the assemblage level across the eleven zoogeographic realms. Lines of best fit are based on a simple linear regression.

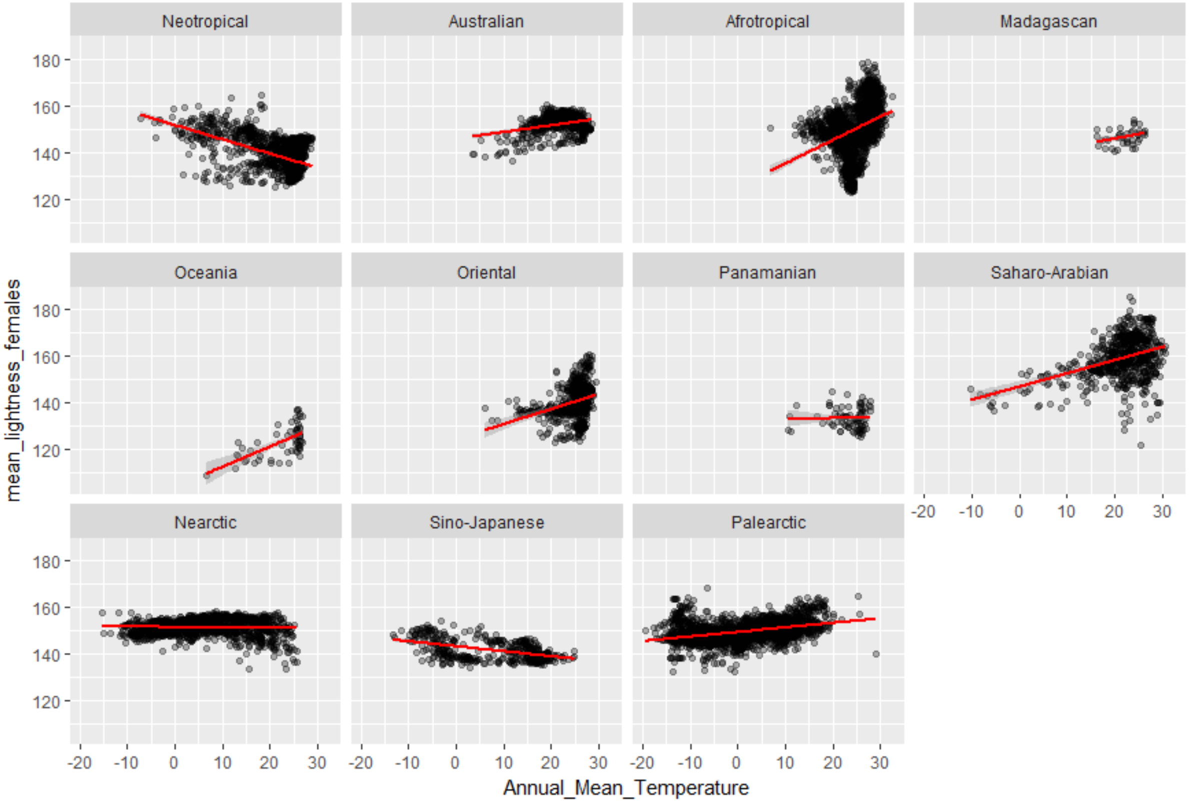

**Fig. S5.** Scatterplots depicting the correlation between annual mean temperature and annual precipitation at the assemblage level across the eleven zoogeographic realms. Lines of best fit are based on a simple linear regression.

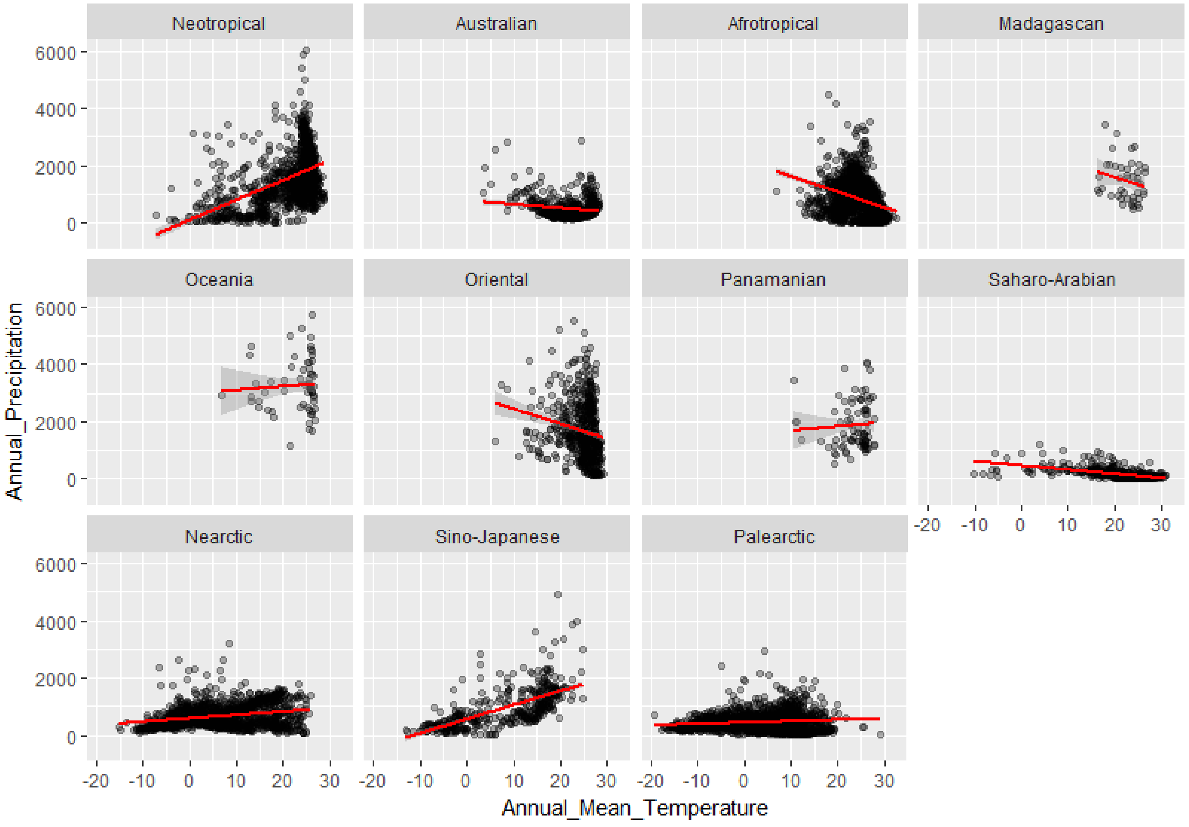

**Supplementary Tables** (see file SupplementaryTables.xlsx)

**Table S1.** Species-level results from comparative analyses examining the effects of mean temperature, annual precipitation and tree cover on male and female plumage lightness across 5809 species of passerine birds. Shown are results from the best model which includes all linear effects and significant quadratic effects, the full model including all effects and the model with only linear effects. In each case, results represents model averaged estimates from 1000 different phylogenies to incorporate phylogenetic uncertainty. For each model we report model averaged estimates, their SEs and associated p-values (median, min and max based on the 1000 different models), proportion of p-values < 0.05 (prop. significant), mean AIC and lambda values.

**Table S2.** Assemblage-level results from linear mixed models examining the effects of mean temperature, annual precipitation and tree cover on average female and male plumage lightness across 9715 assemblages (1° x 1°) of passerine birds. Shown are results from the best model which includes all linear effects and significant quadratic effects, the full model including all effects and the model with only linear effects. All models include a residual autocorrelation vector as a fixed covariate to account for spatial autocorrelation (rac.w) following (Crase *et al.* 2012). Mixed models included zoogeographic realm as random intercept and the interactions between realm and/or temperature, precipitation or tree cover as random slopes.

**Table S3.** Results from meta-regressions assessing how the correlation between annual precipitation and mean temperature affects the relationship between climatic and environmental variables on species-level female and male plumage lightness across passerine families ( N = 65 families with 10 or more species, for results using families with 15 or more species see Table S5). In each case, the dependent variable was the effect (slope) of a single environmental variable (mean temperature, annual precipitation or tree cover) on plumage lightness as obtained from a model including all three variables together or each predictor separately (linear effects only, see text), computed separately for each family. The independent variable was the correlation coefficient between mean temperature and annual precipitation for each family. Meta-regressions were carried out separately for each sex. Meta-analytical means (the model intercepts) were computed for $r_{rain\text{-}temp} = 0$.

**Table S4.** Results from meta-regressions assessing how the correlation between annual precipitation and mean temperature affects the relationship between climatic and environmental variables on assemblage –level female and male plumage lightness across zoogeographic realms (N = 11). In each case, the dependent variable was the effect (slope) of a single environmental variable (mean temperature, annual precipitation or tree cover) on plumage lightness as obtained from a model including all three variables together or each predictor separately (linear effects only, see text), computed separately for each realm. The independent variable was the correlation coefficient between mean temperature and annual precipitation for each realm. Meta-regressions were carried out separately for each sex. Meta-analytical means (the model intercepts) were computed for $r_{rain\text{-}temp} = 0$.

**Table S5**. Results from meta-regressions assessing how the correlation between annual precipitation and mean temperature affects the relationship between climatic and environmental variables on species-level female and male plumage lightness across passerine families (N = 56 families with 15 or more species). In each case, the dependent variable was the effect (slope) of a single environmental variable (mean temperature, annual precipitation or tree cover) on plumage lightness as obtained from a model including all three variables together or each predictor separately (linear effects only, see text), computed separately for each family. The independent variable was the correlation coefficient between mean temperature and annual precipitation for each family. Meta-regressions were carried out separately for each sex. Meta-analytical means (the model intercepts) were computed for rrain-temp = 0.
